# Infant cortical tracking of speech shows emerging spatial release from masking in the first year of life

**DOI:** 10.64898/2026.01.20.700695

**Authors:** Farhin Ahmed, Qianxun Zheng, Jenny Mcllwain, Talat Jabeen, Bonnie K. Lau

## Abstract

Natural listening environments are filled with competing sounds. One mechanism that helps overcome this challenge is spatial release from masking (SRM), whereby spatial separation between a target signal and interfering sounds improves perception of the target. SRM has been observed in a wide range of species including songbirds, crocodilians, ferrets, and human adults, suggesting an evolutionarily conserved strategy for listening in noise. However, an important question remains: Is an infant’s developing brain able to tap into the same mechanism to cope with the noisy world? To address this question, we recorded electroencephalography (EEG) from 7- and 11-month-old infants (N = 53) and adults (N = 20) as they listened to natural speech in quiet, collocated noise, and spatially segregated noise. Our results revealed that both infants and adults showed robust cortical tracking of speech across quiet and noisy listening conditions. For adults, spatial separation between the target speaker and distracting talkers led to enhanced cortical tracking of the target speech, consistent with SRM. Infants also showed SRM, with stronger tracking in segregated than collocated noise, although the effect was confined to a frontal-central region rather than broadly distributed across the scalp as in adults. These findings provide the first neurophysiological evidence that, although still immature, the developing brain can benefit from spatial cues in the first year of life. Together, these results add new insight into how the infant brain solves a fundamental perceptual problem - identifying a relevant voice in a noisy environment - using a mechanism that is evolutionarily grounded.

## INTRODUCTION

Humans live in environments that are inherently noisy. From birth, infants are exposed to a constant stream of competing sounds not only in their homes (Benítez-Barrera et al., 2020), but also in daycares (Frank & Golden, 1999; Golden & Frank, 2000) and on playgrounds (Stadnik, 2021; Wu, 2006). Noise levels in these settings frequently exceed recommended standards for optimal listening and learning (Manlove et al., 2001; Picard & Bradley, 2001). These elevated noise levels continue to persist throughout the first two years of life, a critical period for speech and language development (Kuhl, 2010). As a result, infants must learn to parse relevant speech information while suppressing background noise long before their central auditory systems are fully mature.

One mechanism that supports listening in noisy environments is spatial release from masking (SRM). When competing sound sources are spatially segregated from a target talker, adults show improvement in speech perception (Bronkhorst & Plomp, 1988; Dirks & Wilson, 1969; Freyman et al., 1999). Importantly, SRM is not unique to human listeners. It has been observed across numerous species, including zebra finches, budgerigars, ferrets, and crocodilians (Dent et al., 2009; Hine et al., 1994; Thévenet et al., 2022). The presence of SRM in organisms with vastly different neural architectures implies that leveraging spatial cues to resolve acoustic competition is an evolutionarily conserved strategy employed during complex listening. At the neural level, SRM depends on a cascade of processes operating at multiple stages of the auditory system, including accurate encoding of the incoming sound mixture by the ear and auditory nerve fibers (Bronkhorst, 2015), early brainstem encoding of binaural cues such as interaural time and level differences (Franken et al., 2018; Pecka et al., 2008), and subsequent integration of spatial information in higher auditory regions (van der Heijden et al., 2019). Because many of these processes rely on precise neural timing, mature spatial tuning, and coordinated activity across distributed brain networks, they are likely sensitive to developmental immaturity (Mrsic-Flogel et al., 2003).

This intersection between evolutionary conservation and neural immaturity motivates the present study. Although SRM has been observed across species, the human auditory system undergoes prolonged postnatal development, raising the possibility that this mechanism is not fully functional in infancy. Infancy thus presents a critical stage in which a biologically established strategy for listening in noise must function within an immature neural system. Beyond its theoretical importance, this question also has practical relevance, as limitations in early spatial hearing may reduce access to linguistic input and disrupt early word learning (Evans & Maxwell, 1997; Maxwell & Evans, 2000; McMillan & Saffran, 2016).

Behavioral studies in humans have examined SRM in both adults and children (Ching et al., 2011; Yuen & Yuan, 2014), but its developmental trajectory remains unclear. Some studies report reliable SRM by early childhood (Ching et al., 2011; Garadat & Litovsky, 2007; Litovsky, 2005), or even in toddlers aged 3 years (Lovett et al., 2012), whereas others suggest that spatial hearing continues to mature through adolescence (Cameron et al., 2006; Vaillancourt et al., 2008; Wightman & Kistler, 2005). Despite this work, SRM has not been studied during the first year of life, largely because infants cannot perform the behavioral tasks typically used to assess speech perception in noise. As a result, our understanding of how spatial hearing emerges at the earliest stages of development remains limited.

A robust neural signature of speech processing is cortical speech tracking, a phenomenon in which the low-frequency brain activity tracks the amplitude envelope of continuous speech (Aiken & Picton, 2008; Lalor & Foxe, 2010; Luo & Poeppel, 2007). Importantly, recent work has demonstrated that cortical speech tracking is robust in preverbal infants (Attaheri, Choisdealbha, et al., 2022; Attaheri, Panayiotou, et al., 2022; Jessen et al., 2019; Jessica Tan et al., 2022; Kalashnikova & Burnham, 2018; Menn et al., 2022), including newborns (Ortiz Barajas et al., 2021). However, to date, infant studies have focused on quiet listening conditions, leaving open the question of how noise and spatial cues influence cortical tracking early in life.

We addressed this gap by recording electroencephalography (EEG) from 7- and 11-month-old infants and adults, while they listened to speech in quiet, collocated noise, and spatially segregated noise. Cortical speech tracking was quantified using the multivariate Temporal Response Function (mTRF) approach (Crosse et al., 2016), and the difference in neural tracking between collocated and segregated noise conditions served as a neural index of SRM. This framework enabled objective comparisons of noise effects and SRM between infants and adults under ecologically relevant listening conditions, without requiring behavioral responses. Our results provide the first neural evidence of SRM in infants, indicating that the mechanisms supporting SRM are present, albeit immature, during the first year of life.

## METHODS

### ***A.*** Participants

Participants for this study included 53 infants - 25 7-month-olds (7M; 30.4 ± 1 weeks; 12 female) and 28 11-month-olds (11M; 48.3 ± 1 weeks; 13 female), as well as 20 adults (20.8 ± 1.8 years; 12 female) [mean ± standard deviation (SD)]. Data from 11 additional infants were excluded due to excessive movement or fussiness (n=8), an insufficient number of usable trials (n=2), and hair thickness interfering with EEG signal quality (n=1).

All infants were born full-term, had no history of otitis media within two weeks of testing, passed their newborn hearing screening, and had no history of health or developmental concerns. Parent-reported race and ethnicity included 3 Asian, 23 White, 16 More Than One Race, and 11 Unknown. Infants completed a single testing visit lasting approximately one hour. At the end of each session, a tympanometric screen was conducted to confirm absence of middle-ear fluid.

All adult participants reported clinically normal hearing with no history of health or neurodevelopmental concerns that would interfere with participation. Self-reported race included 11 Asian, 8 White, and 1 Pacific Islander. On the day of testing, a standard audiometric screening (<20 dB HL) at octave frequencies between 500 and 8000 Hz as well as a tympanometric screening was conducted for study inclusion. Data from one additional adult was excluded due to a technical issue.

All procedures were approved by the Institutional Review Board at the University of Washington. Written informed consent was obtained from parents or legal guardians of infant participants while adult participants provided written informed consent for their own participation. All participants received monetary compensation for their time.

### ***B.*** Experimental conditions

Participants completed a passive listening EEG recording involving three conditions: quiet, collocated, and segregated (Fig. 1). In the quiet condition, the target speech was presented alone from a loudspeaker at 0° azimuth. In the collocated condition, the target speech along with a 4-talker babble noise were all presented from the same loudspeaker at 0° azimuth. In the segregated condition, the target speech was again presented from 0° azimuth while two distracting talkers (1 male, 1 female) was presented at -90° azimuth and two (1 male, 1 female) at +90°, creating spatial separation between target and noise. Each condition consisted of 10 1-minute-long trials resulting in a total recording duration of 30 minutes. Condition and trial order were randomized across participants.

**Figure 1:**
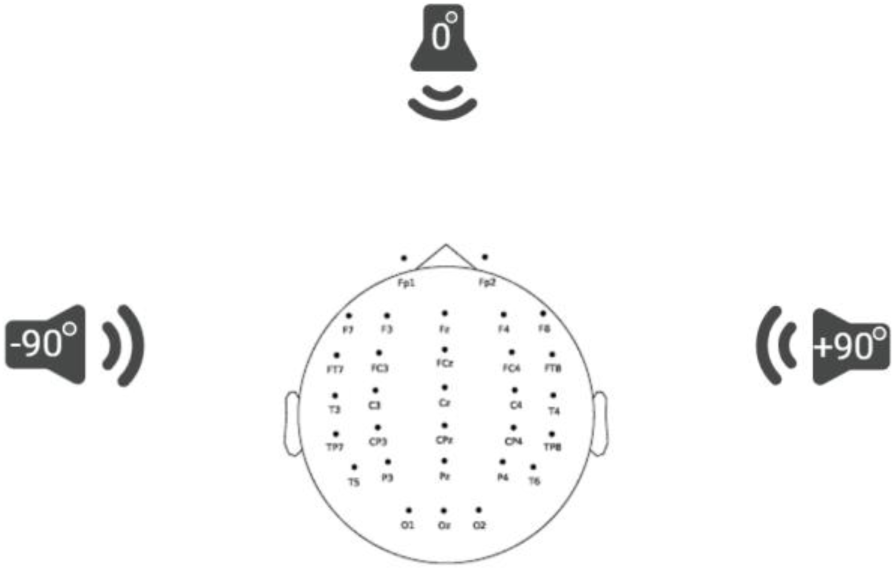
Schematic diagram of loudspeaker arrangement (0°, −90°, +90° azimuth) and the three listening conditions: quiet (only infant-directed target speech at 0°), collocated (target speech and 4-talker babble noise both at 0°), and segregated (target speech at 0° with 2 distracting talkers spatially separated at −90° and 2 distracting talkers at +90°).

### ***C.*** Stimuli

The target stimuli consisted of naturally produced infant-directed speech (IDS). IDS is characterized by a slower rate of speech, longer duration, elevated pitch with a wider pitch range, and simplified grammatical structure (Frühholz & Belin, 2018; Kitamura & Burnham, 2003; Panneton et al., 2006; Payne et al., 2015). Compared to adult-directed speech, IDS elicits stronger cortical tracking in infants (Kalashnikova et al., 2018) and promotes attention to the stimuli (Cooper & Aslin, 1990; Fernald, 1985; Pegg et al., 1992). The IDS stimuli were produced by two female native speakers of English. The recording took place in a sound-attenuated booth with the speaker seated about 25 cm from a microphone. Each speaker produced approximately 30 minutes of IDS, from which two trained listeners manually selected 15 1-minute segments per speaker, yielding a total of 30 minutes of target speech. The noise stimuli were constructed from four audiobooks narrated by American English speakers (two male, two female). All stimuli were inspected for artifacts and then RMS-normalized in Praat (Boersma & Weenink, 2022). Stimuli were presented at an overall level of 70 dB SPL with the signal-to-noise ratio (SNR) between target and distracting talkers set at +5 dB.

### ***D.*** EEG recording

While the stimuli were playing, continuous EEG data were recorded using a 32-channel Biosemi system at a sampling rate of 2048 Hz. During the recording, infants sat on their caregiver’s lap inside a sound-attenuated booth approximately 1m from each loudspeaker. A research assistant was present with them in the booth. The assistant’s primary responsibility was to maintain the infant’s attention, arousal, and behavior to optimize data quality during acquisition. The assistant utilized visually engaging but silent materials, such as brightly colored toys, picture books, and puppets, to hold the infant’s attention. A behavioral coding scheme was also implemented by an experimenter who was live coding for attention and state of the infant throughout the session. Breaks were offered during the session, if needed. If regaining the infant’s attention was not possible, the recording session was ended. All experimenters and research assistants in the laboratory underwent an extensive training protocol prior to assisting with infant data acquisition. On average, infants provided 27.78 ± 4.33 minutes of usable EEG data per session. No significant difference in data duration was observed across the 3 conditions (χ²(2) = 1.86, *p* = 0.39; Friedman test).

Adult participants sat alone in the booth and watched a silent movie. All adult participants completed the full experiment.

### ***E.*** EEG preprocessing

Both infant and adult EEG data were preprocessed identically to ensure consistency and to rule out any group differences that might arise from discrepancies in the preprocessing pipeline. EEG pre-processing was conducted offline with custom MATLAB scripts (The MathWorks, Inc., 2022) incorporating the EEGLAB toolbox (Delorme & Makeig, 2004). EEG data were filtered into 1-4 Hz frequency band based on prior evidence that infants demonstrate stronger cortical speech tracking in the delta band (1–4 Hz) compared to the theta band (4–8 Hz) (Attaheri, Choisdealbha, et al., 2022; Attaheri et al., 2024). Filtering was conducted using zero-phase shift bandpass Hamming windowed FIR filters via the *pop_eegfiltnew* function in the EEGLAB toolbox with a transition band width of 2 Hz and cutoff frequencies at -6 dB. The EEG data was then down sampled to 128 Hz for better computational efficiency. Next, the EEGLAB function *clean_asr* was used to remove transient artifacts including muscle activity, eye blinks, and movement, as in prior infant EEG studies (Attaheri, Choisdealbha, et al., 2022; Attaheri, Panayiotou, et al., 2022; Jessica Tan et al., 2022; Kalashnikova & Burnham, 2018). This function implements Artifact Subspace Reconstruction (ASR), an automated EEG artifact rejection method. ASR first identifies the cleanest portion of the data to compute baseline statistics. Then, a 500 msec long sliding-window subspace decomposition is applied on the whole data to detect subspaces with abnormally high variance relative to the baseline data (threshold = 5 SDs). Channels contributing to these high-variance subspaces are reconstructed using a mixing matrix that was calculated on the clean baseline data. Afterwards, channels with excessive noise that could not be corrected with ASR were identified by calculating the kurtosis of each channel, with the threshold for rejection set to 3 SDs away from the mean using the *pop_rejchan* function in EEGLAB. Spherical interpolation was then performed to interpolate the noisy channels. Finally, the EEG data was re-referenced to the average of the two mastoid channels.

### ***F.*** mTRF Analysis

To quantify how well the target speech is encoded in the brain activity, we used the mTRF approach (Crosse et al., 2016, 2021). This method describes the mapping between different features of the speech stimulus and the corresponding neural response recorded with EEG during listening. Although the mTRF framework can be applied to both low-level acoustic and higher-level linguistic features of speech (Di Liberto et al., 2015), here we focused on acoustic representations only given infants’ immature linguistic abilities. Specifically, we extracted two acoustic features corresponding to the target IDS:

- Envelope: The amplitude envelope of the IDS presented in each trial was extracted by applying the Hilbert transform to the raw speech signal. The envelope feature captures the instantaneous magnitude of the acoustic speech signal.
- Envelope Derivative: The first derivative of the broadband envelope was extracted and then half-wave rectified. The envelope derivative feature represents an approximation of the onsets in acoustics and was shown to contribute to predicting neural response (Daube et al., 2019).

The speech features were down sampled to 128 Hz to match that of the EEG signal. We then fit forward mTRF models to estimate the relationship between the two above-mentioned features and the neural response (EEG signal). A forward mTRF can be described as a filter that linearly transforms a set of speech features, S(t) to the corresponding EEG response, R(t) at each channel over a range of time lags: R(t) = TRF_w_ * S(t) + ε, where TRF_w_ are the weights of the filter at every time lag (reflecting the impulse response of the input-output system), and ε represents the residual response not explained by the model. TRF_w_ is estimated by minimizing the mean square error between the actual neural response R(t), and the response predicted by the linear transformation.

We used the mTRF toolbox (Crosse et al., 2016) which solves TRF_w_ using ridge regression: TRF_w_ = (S^T^S + λI)^-1^S^T^R, where λ is the ridge regression parameter that controls for overfitting, I is the identity matrix, and S is the lagged time series of the stimulus feature matrix S(t). We used a time lag window from -100 ms to 300 ms to fit the mTRF model, which is expected to capture most of the stimulus-related neural response of interest based on previous research (Attaheri, Choisdealbha, et al., 2022; Crosse et al., 2016; Di Liberto et al., 2015; Ding & Simon, 2014). The ridge regression parameter λ is optimized using leave-one-out cross validation approach. That is, for each subject in each condition, we trained TRF_w_ on *n*-1 trials (*n* = number of usable trials) for a wide range of λ values across a logarithmic parameter space, ranging from 10^-3^ to 10^8^, and tested the TRF_w_ on the held-out nth trial. This leave-one-out procedure was repeated *n* times so that each trial served as the test set once. The λ value yielding the highest Pearson’s correlation between the predicted and actual EEG responses, averaged across trials and channels, was selected for each subject. Pearson’s correlation at this optimal λ was reported as EEG prediction accuracy (PA).

### ***G.*** Statistical analysis

To determine whether the TRF models performed better than chance, a non-parametric, permutation-based approach was employed (Combrisson & Jerbi, 2015). In this procedure, the true stimulus-EEG pairing is disrupted by applying a random circular shift to the stimulus. The same cross-validation procedure described above was then conducted to compute PA based on the shuffled data. The process was repeated 100 times for each subject in each condition to generate a chance-level or null distribution of PAs. The average across these 100 iterations was reported as the participant’s null PA. Wilcoxon-signed rank tests were then used to determine whether the PAs were significantly above chance at the group level.

Linear mixed effects (LME) modeling was performed to assess whether PAs differed across the fixed effects of age group (7M, 11M, adults) and condition (quiet, collocated, segregated), with a random intercept included to account for individual variability in PA across participants. Significance of the fixed effects was evaluated using Type III ANOVA with Satterthwaite approximation and post-hoc pairwise comparison was performed using estimated marginal means along with False Discovery Rate (FDR) correction. All LME analyses were conducted in RStudio (Posit Software, PBC, 2025) using *lmerTest* version 3.1.3 and *lme4* version 1.1.36.

## RESULTS

### Infants show robust cortical speech tracking in noise

Based on previous research in infants (Attaheri, Choisdealbha, et al., 2022) and children (Van Hirtum et al., 2023; Vander Ghinst et al., 2019), strongest cortical tracking of speech occurs in the delta (1-4 Hz) frequency band of EEG. Thus, multivariate TRFs were fit to describe the mapping of acoustic speech features to EEG signals in the delta band. Pearson’s correlation (r) between the actual and predicted EEG (averaged across all EEG channels) is reported as PA that quantifies the cortical tracking of speech (Fig. 2). To test the PAs against chance, random permutation statistics or null PAs were obtained for each participant. PAs were significantly greater than chance for both 7- and 11-month-old infants across all listening conditions (Wilcoxon signed-rank tests): quiet (Fig. 2, orange bar; *z* = 3.94, *p* < 0.001, 7M; *z* = 3.78, *p* < 0.001, 11M), collocated (Fig. 2, blue bar; *z* = 2.54, *p* = 0.010, 7M; *z* = 3, *p* = 0.002, 11M), segregated (Fig. 2, green bar; *z* = 2.76, *p* = 0.005, 7M; *z* = 3.21, *p* < 0.001, 11M). Together, these results suggest that infants at both 7 months and 11 months of age reliably track the target speech in both quiet and noisy listening environments (Fig. 2).

**Figure 2:**
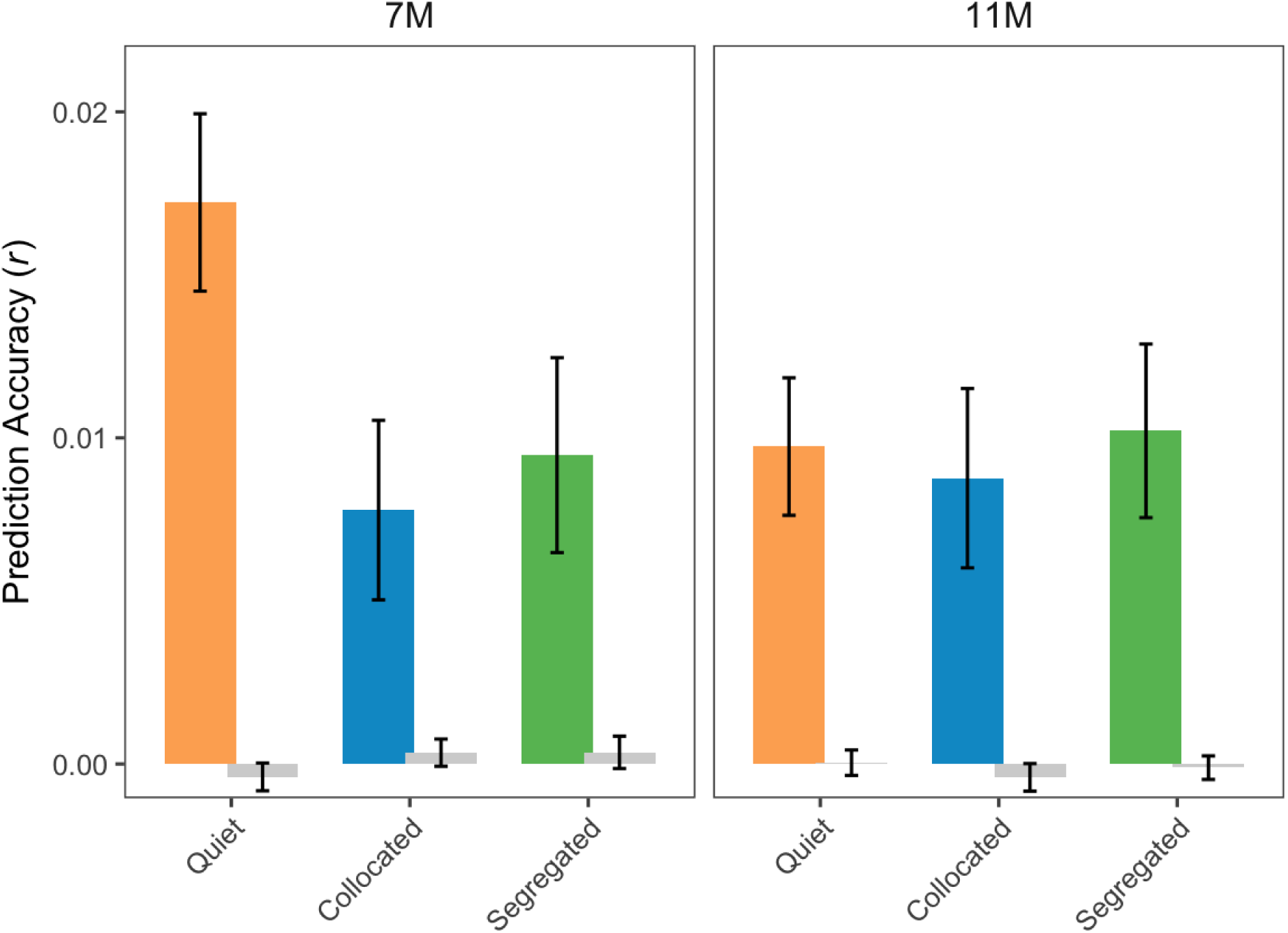
EEG PA in the 1-4 Hz EEG frequency band in all 3 conditions for infants aged 7 months (left) and 11 months (right). Null PAs are shown in gray. PAs are averaged across subjects, all EEG electrodes and trials. Bar plots represent mean ± SEM (standard error of the mean).

To test the effect of age on PA, we fit a LME model. The model revealed no significant main effect of age (*F*(1,152) = 0.78, *p* = 0.38, *η²p* = .005), indicating no difference between 7- and 11-month-olds. Therefore, data from both 3- and 7-month-olds were combined into a single infant group (N = 53) for all subsequent analyses.

### Compared to infants, adults show stronger cortical speech tracking in quiet but are more affected by noise

While infants showed robust tracking of speech in both quiet and noise, we tested adult listeners to serve as a mature comparison group (Fig. 3A). To assess differences in EEG PA across age (infants, adults) and condition (quiet, collocated, segregated), we used a LME model with fixed effects of age, condition, their interaction, and subject-level random intercepts. Indeed, the LME model output revealed a significant main effect of age (Fig. 3; *F*(1, 70.68) = 7.96, *p* = 0.006, *η²p* = 0.1), condition (Fig. 3; *F*(2, 141.19) = 20.05, *p* < 0.001, *η²p* = 0.22), as well as a significant interaction between age and condition (Fig. 3; *F*(2, 141.19) = 9.85, *p* < 0.001, *η²p* = 0.12). This finding suggests that cortical speech tracking differs between infants and adults across listening conditions. As such, post-hoc comparisons were conducted within each age by condition group, with FDR correction applied for multiple comparisons.

**Figure 3:**
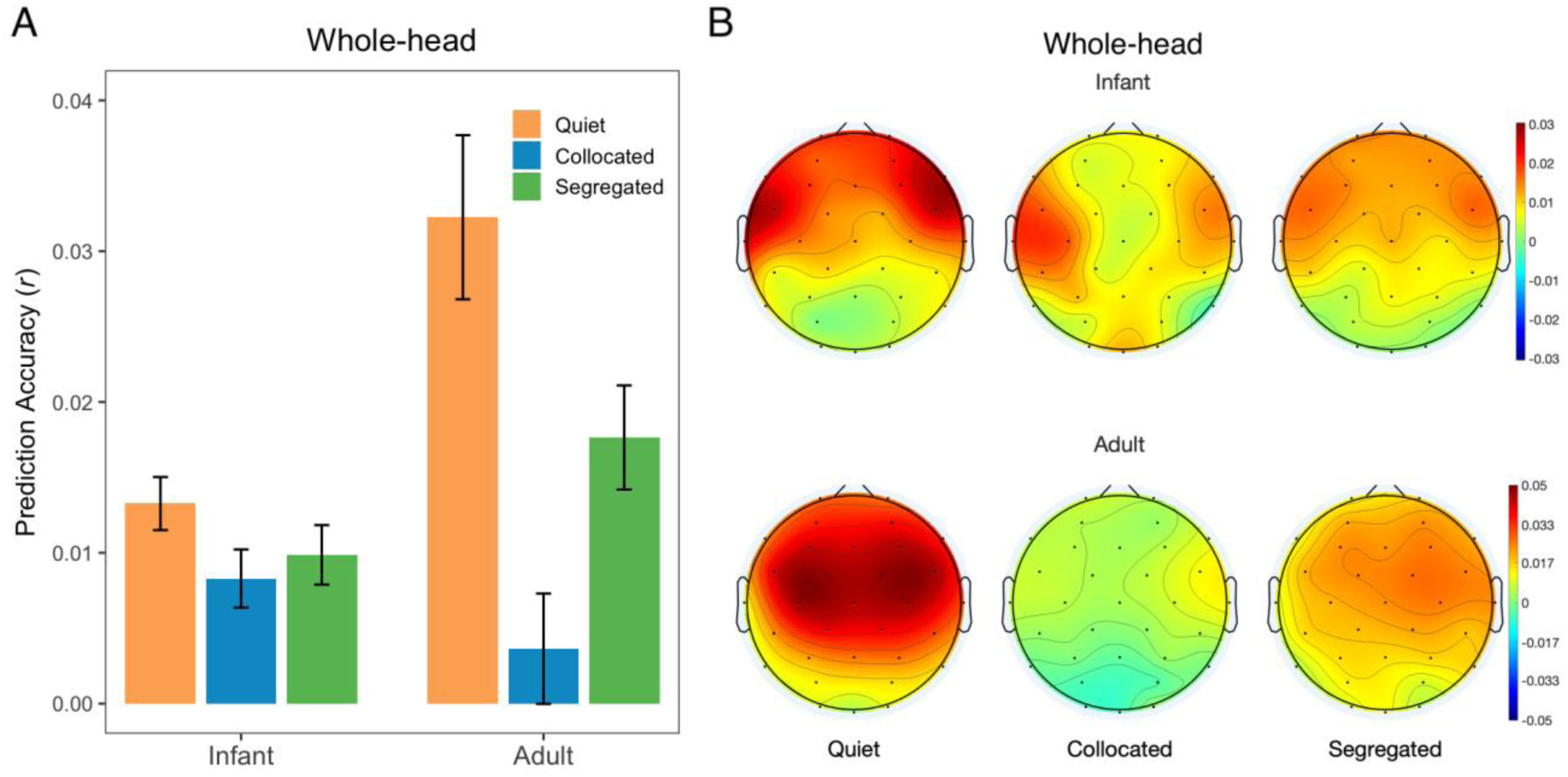
(A) Whole-head prediction accuracies (r) across conditions averaged across all EEG electrodes for infants and adults (mean ± SEM). (B) Topographical distribution of PA across conditions for infants (top) and adults (bottom). Red indicates higher values and blue indicates lower values.

For adults, PA was significantly lower in both collocated and segregated noise compared to quiet (Fig. 3A. quiet vs collocated: *t* = 6.33, *p* < 0.001; quiet vs segregated: *t* = 3.23, *p* = 0.005). This finding suggests that adults’ cortical speech tracking degrades in the presence of noise, regardless of spatial configuration. For infants, we see a similar pattern with the highest mean PAs in the quiet condition and lower PAs in the noise conditions (Fig. 3A, orange vs. blue and green bar). However, this difference did not reach statistical significance across the collocated or segregated noise conditions compared to quiet (quiet vs collocated, quiet vs segregated, all *p* > 0.05). This finding suggests that infants show less effect of noise than adults. Across age groups, adults showed significantly higher PAs compared to infants in the quiet condition (Fig. 3A, orange bar, adults vs infants, quiet: *t* = 4.71, *p* < 0.001) but no group differences were observed in the collocated or segregated noise conditions (Fig. 3A, blue bar, adults vs. infants collocated; Fig. 3A, green bar, adults vs. infants segregated, all *p* > 0.05).

Together, these findings suggest that while adults exhibit stronger cortical tracking in optimal (quiet) listening environments, they are more affected by competing talkers such that their tracking in noisy conditions becomes comparable to that of infants. Infants, on the other hand, show relatively uniform tracking across quiet and noise, potentially reflecting overall immaturity in the neural tracking of speech.

### Infants show SRM localized to fronto-central region

Next, we examined SRM by comparing PAs between segregated and collocated noise conditions. To directly compare the strength of SRM between infants and adults, we computed a neural SRM index as follows:

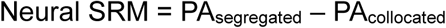

Corresponding topographical maps (Fig. 4A, B) illustrate how this neural SRM index is distributed across electrodes. Adults showed broad, bilateral SRM across the scalp, whereas infants showed SRM focused in a fronto-central region (Fig. 4A top panel; bold black dots represent electrodes where neural SRM is significantly above zero; Wilcoxon signed-rank tests for single electrodes, *p* < 0.05). As such, the largest group-difference in neural SRM appears over centro-occipital scalp region (Fig. 4B bottom panel; bold black dots represent electrodes where group difference in neural SRM is significant; Wilcoxon rank-sum tests, *p* < 0.05). Next, we conducted a region-of-interest (ROI) analysis to quantify SRM within this subset of electrodes (Fig. 4C). Notably, within this ROI, infants exhibited neural SRM comparable to adults with no significant group differences (infants vs adults, *p* > 0.05, Wilcoxon rank-sum test, Fig. 4C). Together, these findings suggest that while adult SRM is broadly distributed across the scalp, infant SRM is limited to fronto-central regions.

**Figure 4:**
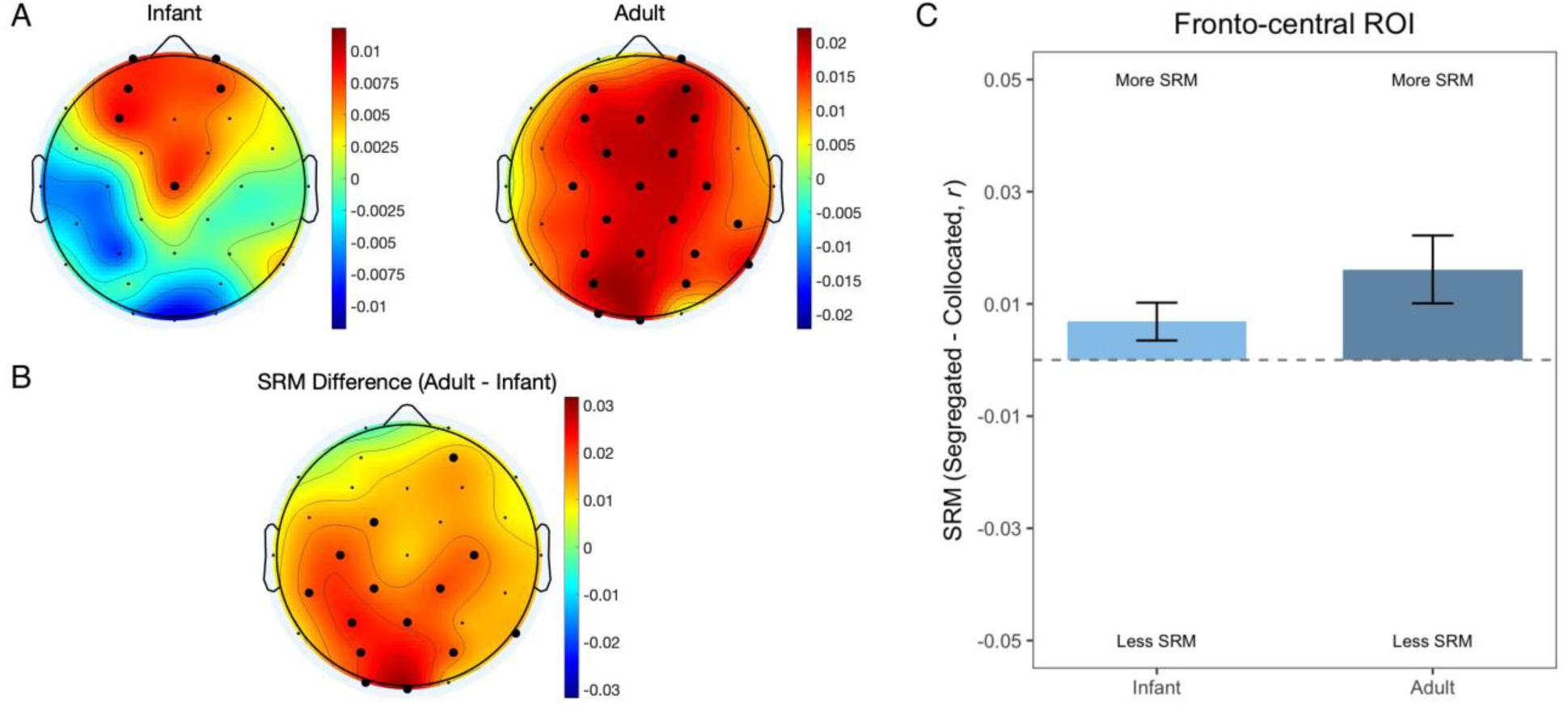
(A) Topographic distribution of neural SRM, where red indicates more SRM and blue indicates less. Bold black dots indicate scalp electrodes where neural SRM is significantly greater than zero across subjects. (B) The topographic distribution of SRM difference between infants and adults, where red indicates a greater difference between infants and adults and blue indicates less. Bold black dots indicate scalp electrodes where adult SRM is significantly greater than infant SRM. (C) Neural SRM in infants and adults (mean ± SEM) averaged across 10 fronto-central EEG electrodes (Fp1, Fp2, AF3, AF4, F3, F4, Fz, Cz, FC1, FC2).

### Comparable TRF morphology with delayed peak latency observed in infants

The regression model weights (i.e., the TRFs) can be interpreted in terms of their spatiotemporal dynamics, analogous to examining event-related potentials (Crosse et al., 2016), but with the important difference that TRFs are derived from modeling natural, continuous speech dynamics. This involves assessing how the TRF weights are distributed across the scalp at different time lags between the ongoing speech and the ongoing EEG signals. For example, TRF at a lag of 100 ms indicates the impact of the speech stimulus at time t on the neural response 100 ms later i.e., at time t + 100 ms. To summarize these dynamics, we grouped the EEG channels into frontal, temporal, occipital as well as left-and right-hemispheric ROIs. Figure 5 depicts the time course of the TRF weights averaged across subjects and across these different ROIs and Figure 6 depicts their spatial distribution across lags from -100 to 400 ms. These visualizations provide insights into the amplitude, latency and scalp topography of the ongoing speech-EEG relationship.

**Figure 5:**
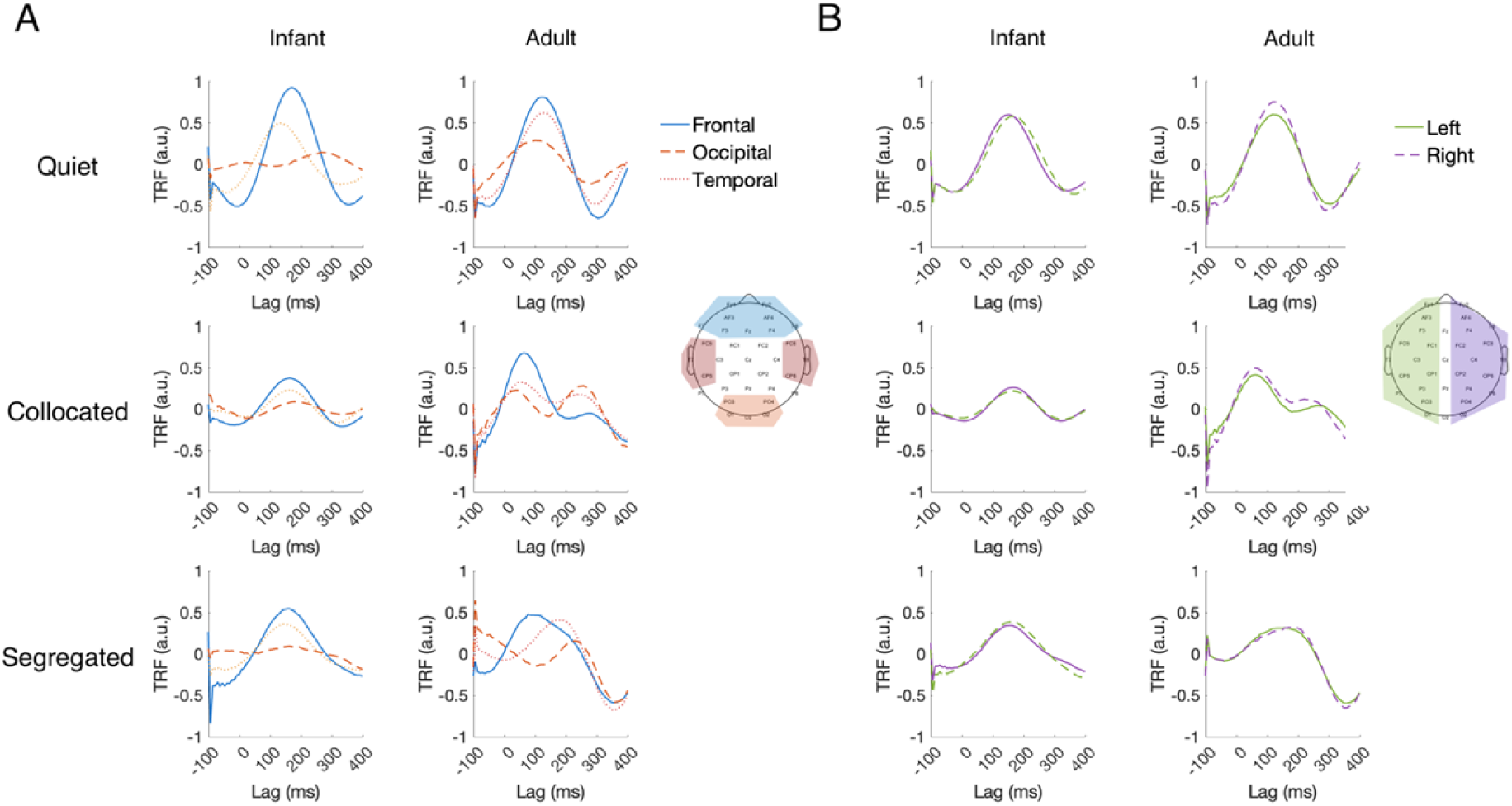
Multivariate TRF weights in infants and adults averaged across subjects (A) at frontal, occipital, and temporal ROIs in infants and adults and (B) at left and right hemispheric ROIs.

**Figure 6:**
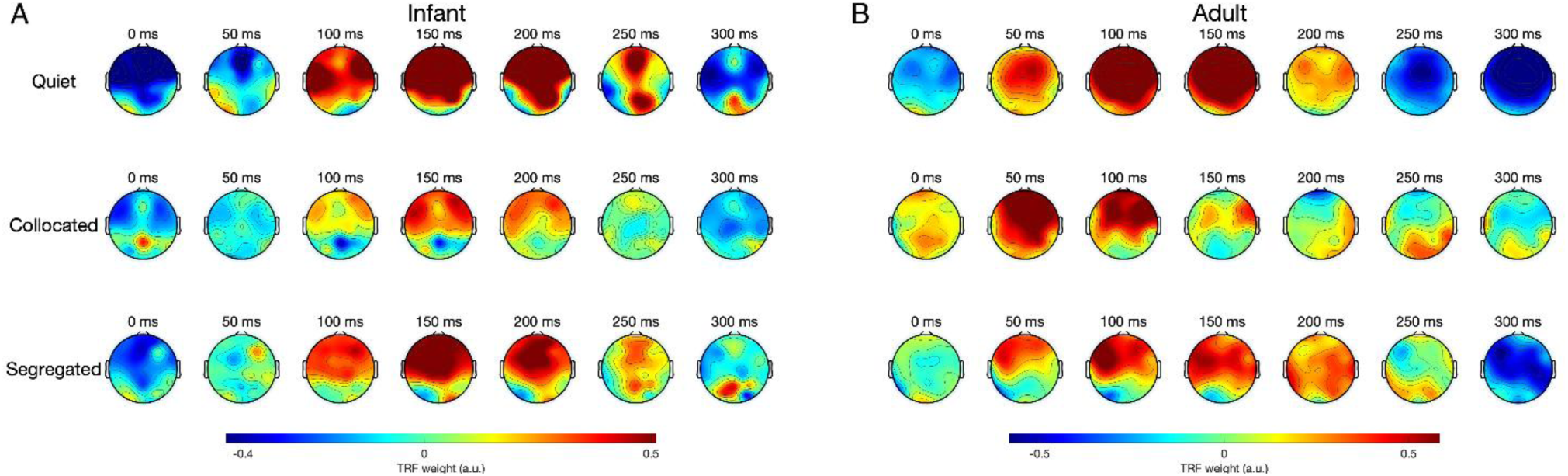
Topographical distribution of TRF weights across different time lags for infants (A) and adults (B). Red indicates higher weights; blue indicates lower weights.

Across both infants and adults, frontal channels showed the strongest TRF weights, followed by temporal channels, while occipital channels showed the lowest weights as expected given that the speech stimulus was presented in the auditory modality rather than visual (Fig. 5A). This is also consistent with standard auditory evoked potentials (AEP) that exhibits larger amplitudes at frontal sites compared with those in occipital areas (Lalor et al., 2009). TRFs from the left and right hemispheric channel sets showed highly similar shapes and amplitudes, indicating no visible hemispheric differences (Fig. 5B).

The temporal profile of the TRFs was broadly similar across conditions with a prominent positive deflection around ∼100 to150 ms and a negative deflection around ∼200 to 300 ms. This temporal profile is consistent with prior EEG studies on cortical tracking of naturalistic speech (Broderick et al., 2018; Crosse et al., 2016; Fiedler et al., 2019; Lalor et al., 2009). Further visual inspection of TRF peak latency revealed a contrast between infants and adults. In general, infant TRF peaks occurred later than adults. In quiet, infants showed longer TRF peak latency (infants: frontal, ∼170 ms; temporal, ∼150 ms; left, ∼150 ms; right, ∼160 ms) than adults (adults: frontal, ∼110 ms, temporal, ∼115 ms, left, ∼110 ms, right, ∼110 ms). A similar pattern was observed in the collocated condition, with infant TRF peaks again occurring markedly later (infants: frontal, ∼175 ms, temporal, ∼175 ms, left, ∼65 ms, right, ∼165 ms) compared to adults (adults: frontal, ∼70 ms, temporal, ∼55 ms, left, ∼60 ms, right, ∼60 ms). In the segregated condition, infant peak TRF latencies fell within a comparable range (infants: frontal, ∼150 ms, temporal, ∼140 ms, left, ∼155 ms, right, ∼160 ms), whereas adult peak latencies were more variable (adults: frontal, ∼95 ms, temporal, ∼165 ms, left, ∼145 ms, right, ∼200 ms).

These group differences in timing were further visualized in the scalp topographies of the TRF weights (Fig. 6). Both infants and adults showed comparable spatial patterns over time characterized by strong fronto-central activity. However, the timing of the spatial distribution was systematically delayed in infants by ∼50-100 ms, consistent with the peak latency shifts observed in Fig. 5. Overall, these spatiotemporal patterns of the TRF weights are similar between infants and adults with a delay in peak latency, suggesting neural immaturity in extracting acoustic information at short timescales of brain activity.

## DISCUSSION

In this study, we investigated how the infant brain tracks continuous speech in noisy, real-world scenarios, with a focus on understanding whether spatial cues provide support for speech processing early in life. Using EEG and TRF modeling, we presented naturally produced IDS in quiet, collocated, and segregated noise. Our results provide the first evidence of cortical speech tracking in noise in pre-verbal infants, extending previous infant studies limited to listening in quiet. Comparison between infants and adults further revealed that while adults exhibit robust neural signature of SRM across the scalp, infants showed SRM focused in a front-central region, suggesting that spatial benefits are evident but still maturing during the first year of life. Finally, infants showed delayed TRF peak latencies relative to adults, indicating slower underlying cortical dynamics during speech tracking in both quiet and noise.

Cortical tracking of speech has been extensively studied in adults, particularly using the TRF approach with naturalistic, non-repetitive stimuli such as movies (Whittingstall et al., 2010) or conversational speech (Broderick et al., 2018; Di Liberto et al., 2015; Ding & Simon, 2012; Fiedler et al., 2019; Puvvada & Simon, 2017), which offer high ecological validity and engage listeners more deeply than isolated, artificial stimuli. More recently, studies have shown that TRFs can capture how infants process speech, however, these studies have presented IDS (Kalashnikova et al., 2018), nursery rhymes (Attaheri, Choisdealbha, et al., 2022; Di Liberto et al., 2023), or cartoons (Jessen et al., 2019; Jessica Tan et al., 2022) almost exclusively in favorable quiet listening condition. Yet infants, like adults, encounter complex acoustic environments in everyday life at home, in daycares, and in other settings that contain overlapping talkers. Whether the infant brain can reliably track desired speech under such noisy conditions has remained unknown. To our knowledge, the present study provides the first evidence that the infant brain can indeed track continuous IDS in noise.

We did not observe any difference in delta-band EEG PA in response to acoustic envelope between 7- and 11-month-old infants (Fig. 2). This finding aligns with Attaheri et al. (2022), who used a similar TRF approach (though employing a backward decoding/envelope reconstruction model), and likewise reported no delta-band tracking differences between 7 and 11 months, however, with the strongest tracking emerging at 4 months. Similarly, Di Liberto et al. (2023) reported no age-related changes in acoustic spectrogram tracking between 7- and 11-month-olds, while again observing the strongest tracking in 4-month-olds. Together, these findings point towards the possibility that the basic acoustic tracking in quiet may develop rapidly from birth to early infancy and then plateau as the brain becomes increasingly sensitive to phonetic, prosodic, and other higher-level linguistic features. Future longitudinal study with larger cohorts will be crucial for clarifying this developmental trajectory.

An interesting finding in our study is that infants did not show differences in cortical tracking of IDS (target) between quiet and either of the noise conditions, even though adults showed the expected degradation in noise. One possibility is that this pattern reflects early developmental stage of auditory scene analysis. Adults possess mature top-down mechanisms for parsing competing streams, enabling them to distinguish the target from distractors. However, this selective processing requires the target to compete with the background, and that competition reduces target tracking in noise relative to quiet. In contrast, infants cannot (or not yet strongly) distinguish target from distractor, resulting in little neural competition. As a result, cortical responses in quiet and noise appear similar, reflecting a more global tracking of the overall acoustic mixture with limited differentiation between target and distracting talkers. This interpretation aligns with behavioral studies showing that infants up to 7–9 months of age often fail to detect a tone embedded in broadband noise (Werner, 2007) – a result often attributed to reduced contrast between target and noise rather than an inability to hear the tone per se (Kushnerenko et al., 2013; Werner & Boike, 2001). Along the same line, our EEG results suggest that infants may treat elements of the auditory scene more uniformly, leading to comparable cortical tracking in quiet and noise. Importantly, while the behavioral work examined non-speech stimuli, our findings extend this pattern to natural speech using an EEG-based measure.

The primary question addressed in this study is how the ability to use spatial cues i.e., SRM emerges early in life to improve listening under noisy conditions. Adults in our sample showed a clear SRM benefit: their tracking of target speech was significantly stronger when the target and distracting talkers were spatially segregated compared with collocated. For infants, SRM was observed over a number of fronto-central electrodes, and the adult vs infant difference map revealed the largest group difference over centro-parietal and centro-occipital regions. Together, these observations indicate that while infants do not yet exhibit fully mature SRM across the whole scalp, SRM is evident and localized over fronto-central region. The fronto-central localization of infant SRM is not surprising given that scalp-recorded auditory responses are typically dominant over frontro-central electrodes (Lalor et al., 2009; Näätänen et al., 1982; Picton et al., 1974). Since SRM depends on how well the auditory representation of the target is segregated from competing sounds, it is therefore plausible that SRM-related neural effects first emerge within this canonical auditory processing network and only later generalize across broader cortical networks with development. We therefore interpret our finding as evidence that the neural mechanisms supporting SRM are not yet fully developed in infancy, emerging first in auditory areas engaged in the acoustic encoding of speech.

Evidence from animal models provides an important framework for interpreting this pattern of immature but emerging SRM in human infants. A large body of animal work shows that neural circuits supporting spatial hearing—particularly those encoding interaural time and level differences—undergo extensive postnatal refinement that are shaped by sensory experience. In the auditory brainstem, neurons in the medial superior olive, which are critical for ITD processing, achieve their high temporal precision through refinements in dendritic morphology, cell volume, synaptic excitatory and inhibitory inputs, and membrane electrical potentials after hearing onset (Chirila et al., 2007; Franzen et al., 2020; Rautenberg et al., 2009). Similar neuronal refinements have been reported in the lateral superior olive responsible for interaural level difference processing (Kandler & Gillespie, 2005). Viewed in this context, the absence of a robust, scalp-wide neural SRM in infants, alongside a localized fronto-central signature of emerging SRM, aligns with animal evidence that spatial hearing mechanisms develop gradually and refined through postnatal auditory experience. Our results therefore extend this cross-species framework to humans.

One question that often arises is how reliably the TRF models capture stimulus-related variance in EEG, particularly given the inherently noisy nature of infant recordings. Beyond the permutation-based significance tests (which were robust across groups; Fig. 2), we examined the spatiotemporal structure of the TRF weights themselves to assess whether their morphology aligns with known auditory response dynamics. Across both infants and adults, the TRFs showed the expected pattern reported in prior natural speech EEG studies: weights were near zero at pre-stimulus lags, exhibited a prominent peak between ∼100–150 ms, and returned toward baseline by ∼300 ms (Figs. 5–6), consistent with canonical TRF responses observed over 0–400 ms time lags (Lalor & Foxe, 2010). These visualizations therefore provide converging evidence that the models are capturing meaningful auditory encoding beyond noise. Additionally, TRF amplitudes were broadly similar over left- and right-hemisphere electrodes in both age groups (Fig. 5B). This symmetry aligns with adult TRF literature, which often reports comparable morphology across hemispheres even when lateralization emerges in downstream measures such as PA or attention decoding (Crosse et al., 2016; Fiedler et al., 2019; O’Sullivan et al., 2015). However, this pattern in infants contrasts with one prior infant TRF study reporting larger left-hemisphere responses (Kalashnikova et al., 2018). Several methodological differences may explain this discrepancy: they used only frontal left/right channels, whereas our analyses considered a broader hemispheric distribution; substantially smaller sample size (N = 12 vs. N = 53 in the current study); and they modeled EEG responses in the 1–8 Hz band, while our model focused on 1–4 Hz. Finally, we found that infant TRF peak latencies were generally longer than those of adults. This latency difference is consistent with the idea that speech processing relies on coordinated activity across distributed neural pathways (Hickok & Poeppel, 2007). In infants, these pathways are still developing, leading to slower integration and communication across auditory and higher-order cortical regions, which might be reflected in the longer TRF latencies we observed.

Finally, beyond contributing to a growing body of literature characterizing neural response to natural speech in development, our findings have practical implications. Our approach provides objective neural measure of speech-in-noise and spatial hearing for infants that do not rely on non-sensory factors such as attention or task compliance, complementing studies utilizing behavioral methods such as head-turn or preferential listening (Hall et al., 2004; Werner, 2007; Werner & Boike, 2001). Such neural metrics could eventually be incorporated into clinical batteries to support early identification of listening difficulties in infancy. Building on this implication, several future directions stem from this work. A natural next step is to follow infants longitudinally to track how emerging SRM matures during toddlerhood and early childhood. In parallel, it will be important to examine when the longer TRF latencies observed here in infants catch up with that of adults. Importantly, delayed TRF latencies have been reported in adults with hearing loss (Gillis et al., 2022) and in children with cochlear implants (Federici et al., 2025), suggesting that TRF timing may be a sensitive developmental marker worth following closely. Additionally, the TRF framework can be extended to higher-order speech features such as phonetic, lexical, prosodic, and semantic representations (Broderick et al., 2018, 2018; Di Liberto et al., 2015; Heilbron et al., 2022; Teoh et al., 2019). Indeed, recent evidence has indicated that phonetic encoding emerges and progressively strengthens in the infant brain throughout the first year of life (Di Liberto et al., 2023). As a future direction, it would be valuable to investigate how noise and spatial cues affect these levels of processing. Importantly, brief unilateral conductive hearing loss introduced in a mouse model during critical developmental periods disrupted cortical sensitivity to interaural cues (Polley et al., 2013). Future studies could therefore examine whether congenital or early-onset hearing loss similarly alters the development of SRM in human infants. Finally, linking our neural signature of SRM to behavioral outcomes, such as language acquisition would be an important translational step toward better understanding and ultimately supporting real-world communication in early childhood.

## Acknowledgements

This research was supported by R01 DC022585 and R00 DC016640 to BKL, T32DC005361-21 to F.A. and T.J. For Open Access, the authors have applied for a CC BY public copyright license to any Author Accepted Manuscript version. Finally, we thank the participants and their families for their contributions to this research.

## Data availability

Data are available from the corresponding author upon request.

